# Magnetic Stimulation Allows Focal Activation of the Mouse Cochlea

**DOI:** 10.1101/2022.01.11.475825

**Authors:** Jae-Ik Lee, Richard Seist, Stephen McInturff, Daniel J. Lee, M. Christian Brown, Konstantina M. Stankovic, Shelley Fried

## Abstract

Cochlear implants (CIs) strive to restore hearing to those with severe to profound hearing loss by artificially stimulating the auditory nerve. While most CI users can understand speech in a quiet environment, hearing that utilizes complex neural coding (e.g., appreciating music) has proved elusive, probably because of the inability of CIs to create narrow regions of spectral activation. Several novel approaches have recently shown promise for improving spatial selectivity, but substantial design differences from conventional CIs will necessitate much additional safety testing before clinical viability is established. Outside the cochlea, magnetic stimulation from small coils (micro-coils) has been shown to confine activation more narrowly than that from conventional micro-electrodes, raising the possibility that coil-based stimulation of the cochlea could improve the spectral resolution of CIs. To explore this, we delivered magnetic stimulation from micro-coils to multiple locations of the cochlea and measured the spread of activation utilizing a multi-electrode array inserted into the inferior colliculus; responses to magnetic stimulation were compared to analogous experiments with conventional micro-electrodes as well as to the responses to auditory monotones. Encouragingly, the extent of activation with micro-coils was ∼60% narrower than that from electric stimulation and largely similar to the spread arising from acoustic stimulation. The dynamic range of coils was more than three times larger than that of electrodes, further supporting a smaller spread of activation. While much additional testing is required, these results support the notion that coil-based CIs can produce a larger number of independent spectral channels and may therefore improve functional performance. Further, because coil-based devices are structurally similar to existing CIs, fewer impediments to clinical translational are likely to arise.

## INTRODUCTION

More than 430 million people worldwide, ∼5% of the world’s population, live with disabling hearing loss, making it the most common sensory deficit(1). The World Health Organization (WHO) estimates that this number will grow to 700 million by 2050(1). There are significant associations between hearing impairment and reduced quality of life, increased risk for dementia, and/or an inability to function independently(2). In the most common type of hearing loss, called sensorineural hearing loss (SNHL), there is often a loss of the sensory hair cells that transduce sound-induced vibrations within the cochlea into neural activity; this loss precludes the use of sound-amplifying hearing aids as a potential treatment. Instead, a cochlear implant (CI) can be implanted to electrically stimulate spiral ganglion neurons (SGNs), the neurons downstream from hair cells. CIs are generally effective for enabling speech comprehension, but typically to only about 65% of normal(3-6). In addition, there is a significant reduction in CI performance when background noise levels are high and most CI users also cannot appreciate music(7, 8). Thus, despite the unquestionable benefit provided by existing devices, there is room for improvement.

The limitations in performance are thought to arise largely from the small number of independent spectral channels created by CIs. In contrast to the large number of independent channels arising from the ∼30,000 SGNs in the intact human cochlea, CIs produce as few as 8-10 independent spectral channels(7, 9-11). This is smaller than the number of stimulating electrodes in most existing devices(7, 9). The discrepancy is thought to arise from several intrinsic limitations associated with electric stimulation of the cochlea. For example, the high conductivity of the perilymph within the scala tympani leads to an expansive spread of current in the longitudinal direction (along the tonotopic axis). In addition, the high electrical resistance of the bony wall separating the scala tympani from targeted SGNs within the organ of Corti necessitates an increase in the amplitude of stimulation that results in an even wider spread of activation. Excessive spread from individual electrodes can result in overlap of fields from neighboring electrodes, thereby reducing spectral specificity.

Several novel approaches are under consideration to increase the number of independent channels created by CIs. For example, the genetic insertion of light-sensitive ion channels into SGNs allows their activation to be controlled by light instead of electric fields, resulting in narrower channels(12, 13). Another approach uses electrode arrays that penetrate directly into the auditory nerve (14-16); the reduced separation between electrodes and targeted neurons, along with the elimination of the high resistance barrier, enables better control of activation of the central axons of SGNs. While considerable progress has been made with both approaches, the use of genetic manipulations and/or new surgical techniques raise a number of important safety concerns that will need to be addressed prior to large-scale clinical implementation.

Recent studies have shown that magnetic stimulation from small, implantable coils, referred to as micro-coils, can effectively drive neurons of the CNS while confining activation to a narrow region around each coil(17-20). A coil-based approach is potentially attractive for CIs because magnetic fields are highly permeable to most biological materials, e.g., the bony walls of the scala tympani, and thus activation of SGN processes would not require the same increase in stimulation amplitude required for electric stimulation. Further, the spread of magnetic fields in the scala tympani is less sensitive to the high conductivity of perilymph, further helping to confine activation. While the strength of the fields induced by micro-coils is small, computational studies suggest that the spatial gradient of the resulting fields is suprathreshold (17-22), and simulations specific to the cochlea suggest a multi-turn spiral coil should produce fields strong enough to activate SGNs (23). It is not clear however whether spiral coil designs are best for use in a high-count, multi-coil array designed for the cochlea, as they might reduce the flexibility of the implant and thus could increase the risk for iatrogenic trauma during insertion. Instead, simple bends in micro-wires, recently shown to be effective for the activation of CNS neurons (17-20), may offer an attractive alternative to multi-turn spiral coil designs because they allow coils sizes to be minimized, and thus the flexibility and overall structure of coil-based CIs can be made to match existing implants, thereby reducing a barrier to implementation.

Here, we investigate the ability of magnetic stimulation from bent-wire micro-coils to drive the auditory pathway. We evaluated the efficacy of stimulation and the resulting spectral spread of activation by recording with a multi-electrode array positioned along the tonotopic axis of the inferior colliculus (IC), an auditory nucleus downstream from the cochlea. We show that bent-wire micro-coils can indeed drive auditory circuits effectively and further, that the resulting activation from single micro-coils is significantly narrower than that from traditional electrodes, i.e., approaching the relatively narrow spread produced by acoustic stimuli. Control experiments verified that responses from coils were indeed magnetic in origin and that they did not arise from activation of hair cells. Taken together, our results suggest that further investigation of coil-based CIs is warranted as they may produce a larger number of independent spectral channels than electrode-based CIs and thus could lead to improved clinical outcomes.

## RESULTS

### Responses to acoustic stimulation in hearing animals

A 16-channel recording array was implanted along the tonotopic axis of the contralateral inferior colliculus (IC) in anesthetized mice and used to measure responses to (ipsilateral) acoustic, electric and magnetic stimulation of the cochlea (Figs. 1A and B; MATERIALS AND METHODS). Acoustic stimuli consisted of a series of single frequencies ranging from 8 to 48 kHz, chosen to cover much of the tonotopic range represented in the central division of the mouse IC(24). Multi-unit activity (MUA) recorded from each of the sites in the IC was quantified by analog representation, and the cumulative discrimination index, *d′* (d-prime), was calculated to construct spatial tuning curves (STCs; MATERIALS AND METHODS). Typical STCs of the responses to acoustic stimulation were narrow; examples for 8, 16, and 32 kHz are shown in Figs. 1D-F, respectively. At *d′* equal to 1, the channels with the lowest threshold (‘best’ site, BS) were 12, 9, and 5, respectively (indicated by white stars), which is consistent with the known tonotopic organization of the mouse IC(24). The data for all animals and frequencies tested also show this tonotopic organization (Fig. 1G).

**Fig. 1.**
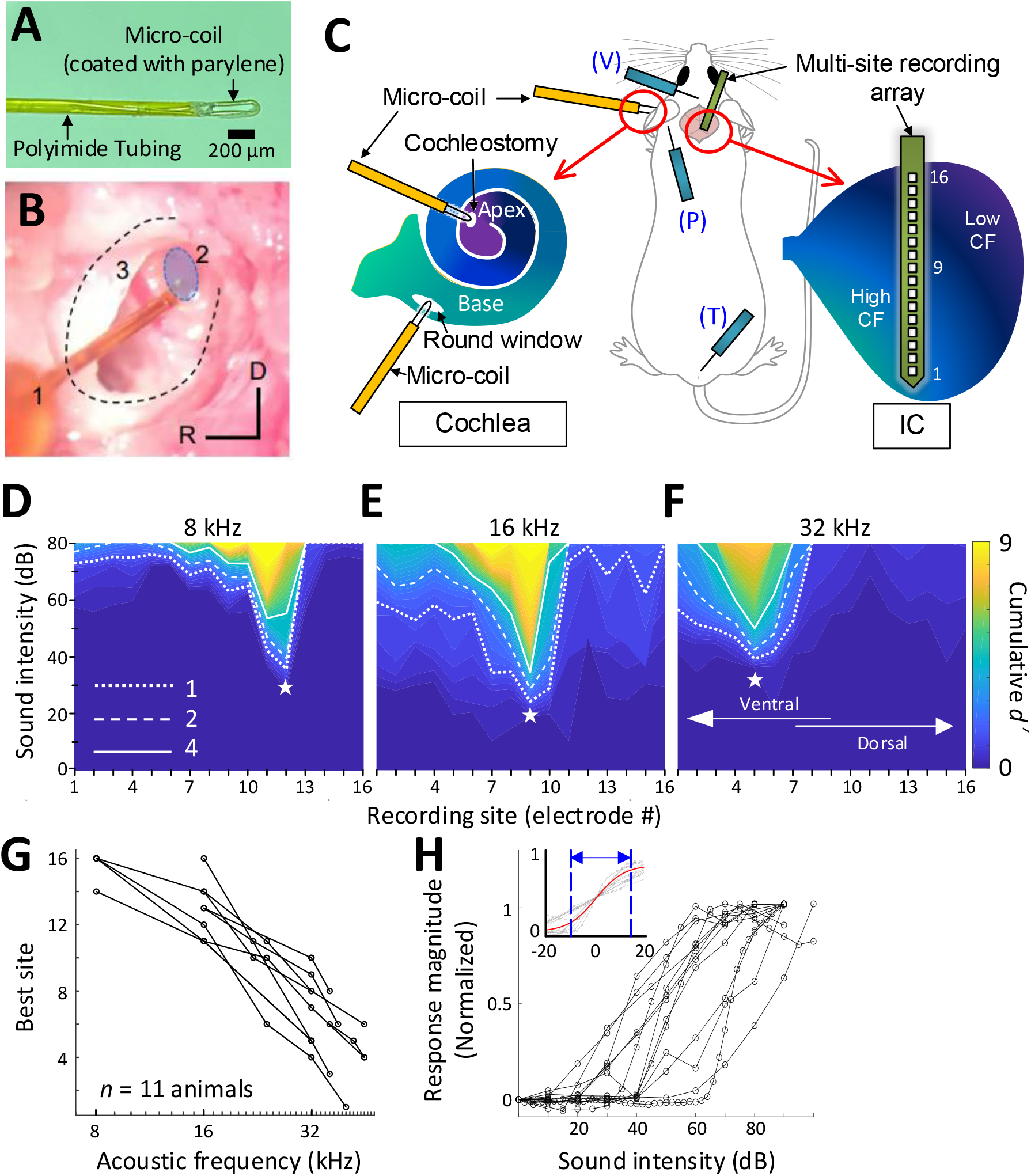
Response to acoustic stimulation measured in the inferior colliculus (IC) **(A)** Photograph of the tip of the micro-coil used in experiments. **(B)** Photograph of the microcoil (1) inserted through the round window of the left cochlea into the basal turn (2, shaded blue). The stapedial artery (3) is visible. The outline of the cochlea is approximated by dashed lines. Axes: R: rostral, D: Dorsal. **(C)** Schematic of the experimental setup depicting the microcoil (orange) inserted into the cochlea (basal and apical turns), the multi-site recording array (green) inserted across the tonotopic axis of the inferior colliculus, and placement of three sub-dermal recording electrodes (blue) into the vertex (V), pinna (P), and tail (T). **(D-F)** Typical spatial tuning curves (STC) of the IC responses to acoustic stimulation (8, 16, and 32 kHz, respectively) recorded from the 16-channel probe positioned in the IC. Response magnitude was quantified with d-prime analysis (see METHODS). The recording site number (x-axis) increases from the IC’s ventral to the dorsal end (low to high characteristic frequency). The recording electrode with the lowest threshold (best site, BS) is marked with a white star. Dotted, dashed and solid lines correspond to cumulative d′ levels of 1, 2, and 4. **(G)** BS for acoustic stimulation from 8 – 48 kHz; lines connect all data from single animals (n = 11). This mapping is used to assign a “characteristic frequency” to each electrode. **(H)** Rate-level functions at BS to 32 kHz normalized to peak rate; individual lines are averaged response from individual animals. Inset plots the same data but normalized such that 50% of the amplitude level that elicited the peak response was assigned the level of 0 dB; the solid red line shows the best-fit sigmoidal curve to all data points.

Consistent with previous studies, increases in supra-threshold sound pressure levels (SPL) typically led to increases in the magnitude of the IC response before saturating at higher intensity levels (Fig. 1H). The dynamic range (DR), defined as the range of stimulus amplitudes for which response strength was between 10% and 90% of the maximum response at BS, averaged 25.96 ± 9.17 dB, consistent with previous reports in mice(25-27). To facilitate comparison of DRs across experiments, especially subsequent responses to magnetic and electric stimulation, the stimulus amplitude that elicited 50% of the maximum response was normalized to a level of 0 dB and the plots of response magnitude vs. stimulus level for individual animals were overlaid (Fig. 1H inset, gray lines). This provides a visual representation of DRs across a given modality; the solid red line is the best-fit sigmoidal curve to all data points, and the dotted lines indicate the 10- and 90-% levels, providing a measure of the DR for the visual overlay of the inset.

### Robust activation of the auditory system by magnetic stimulation

After recording responses to acoustic stimulation, the cochlea was surgically exposed and lesioned to prevent the possibility of acoustic responses arising from magnetic or electric stimulation. For lesioning, distilled water was injected through the round window membrane (see MATERIALS AND METHODS) to induce an osmotic shock to the hair cells(28). Recordings of auditory brainstem responses (ABRs)(29, 30) in mice following intracochlear water instillation demonstrated minimal or no responses up to 75 dB SPL or higher (Fig. 2B).

**Fig. 2.**
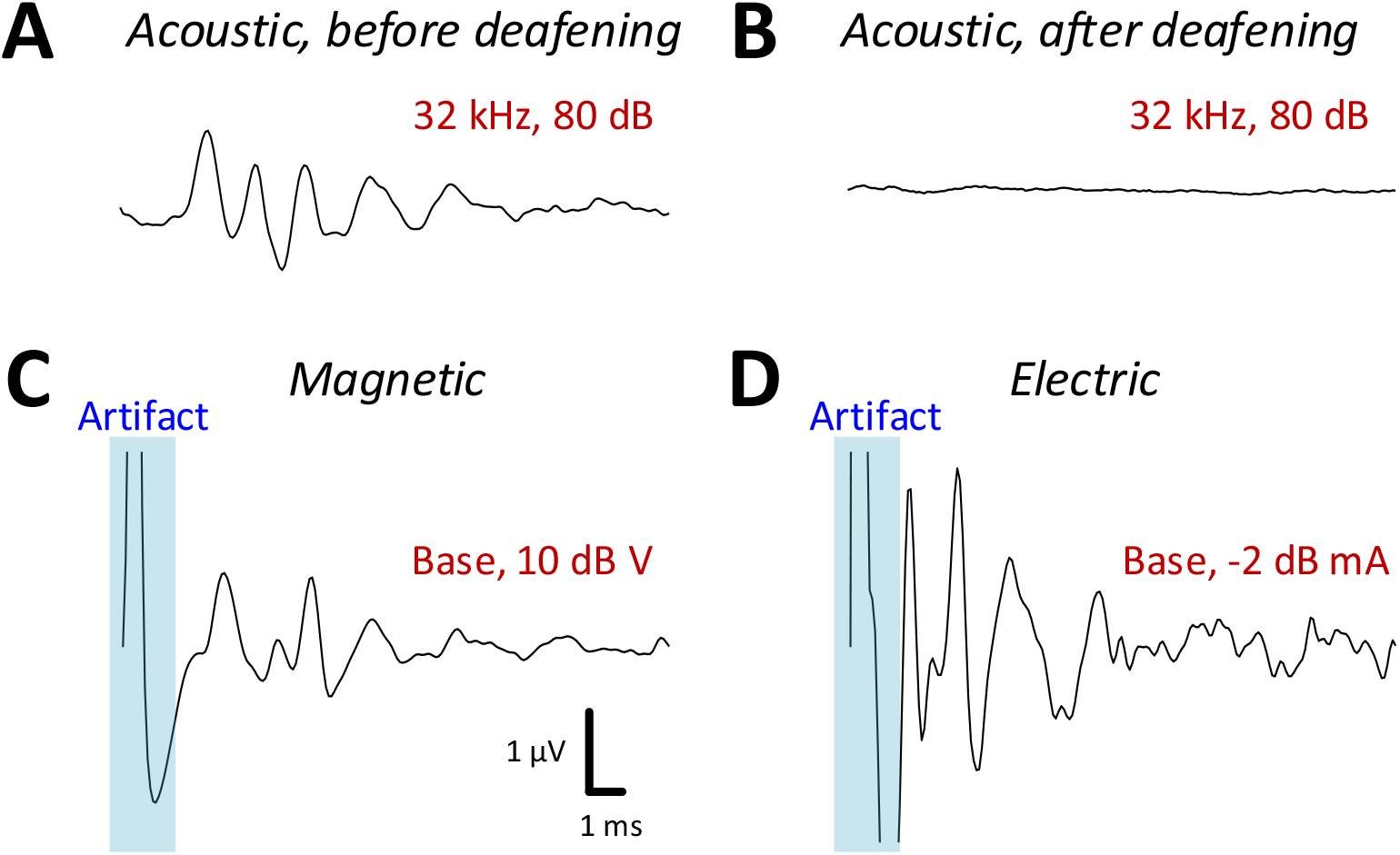
Auditory brainstem responses (ABRs) in response to acoustic, magnetic and electric stimulation. **(A and B)** ABR responses to a 32 kHz monotone; A: control, B: after DI water was injected into the cochlear through the round window. **(C and D)** ABRs from magnetic (C) and electric (D) stimulation (post-deafening). The blue shaded regions identify the portion of the recording obscured by the stimulus artifact.

Following confirmation of severe to profound sensorineural hearing loss on ABR testing, we measured IC responses to both magnetic or electric stimulation, delivered to both basal and apical cochlear. Prior to capturing IC responses, ABRs were recorded each time a coil or electrode was inserted (or reinserted) into the cochlea (see MATERIALS AND METHODS) to provide a relatively quick validation that a given surgical procedure had not damaged the early auditory pathways or the implant itself. ABRs to electric stimulation (eABRs; Fig. 2D) were generally similar to those reported previously in mice(31) and other laboratory animals(32), with multiple peaks occurring within the first few milliseconds following stimulus onset. ABR waveforms to magnetic stimulation (mABR; Fig. 2C) also consisted of a number of peaks, although the amplitudes of the early and later peaks were about the same. The overall appearance of mABRs was closer to that of acoustically evoked ABRs (aABRs) vs. eABRs, although we did not attempt to quantify this observation. Regardless, the generation of robust ABRs to magnetic stimulation strongly suggested that micro-coils can indeed drive the early auditory pathways, and therefore, we proceeded to collect responses from the IC.

Consistent with the presence of robust mABRs, magnetic stimulation also elicited robust neural activity in the IC (Fig. 3A and B). The typical raw response to each modality is shown in Supplementary Fig. 1. Responses to magnetic stimulation were consistent with the tonotopic organization of the cochlea. Responses to stimulation of the basal turn were strong in the ventral portion of the IC, the region known to process high frequencies, with little or no responses observed outside this region (Fig. 3A). Across animals (n = 6), the average characteristic frequency for BSs (extrapolated from Fig. 1G) in response to micro-coil based stimulation of the basal turn was 37.36 ± 4.00 kHz. In contrast, magnetic stimulation of the apical turn elicited only a narrow portion in the dorsal portion of the IC (Fig. 3B), known to process lower sound frequencies. Averaging across the population, the characteristic frequency of the BSs for magnetic stimulation of the apical turn was 8.44 ± 6.58 kHz.

**Fig. 3.**
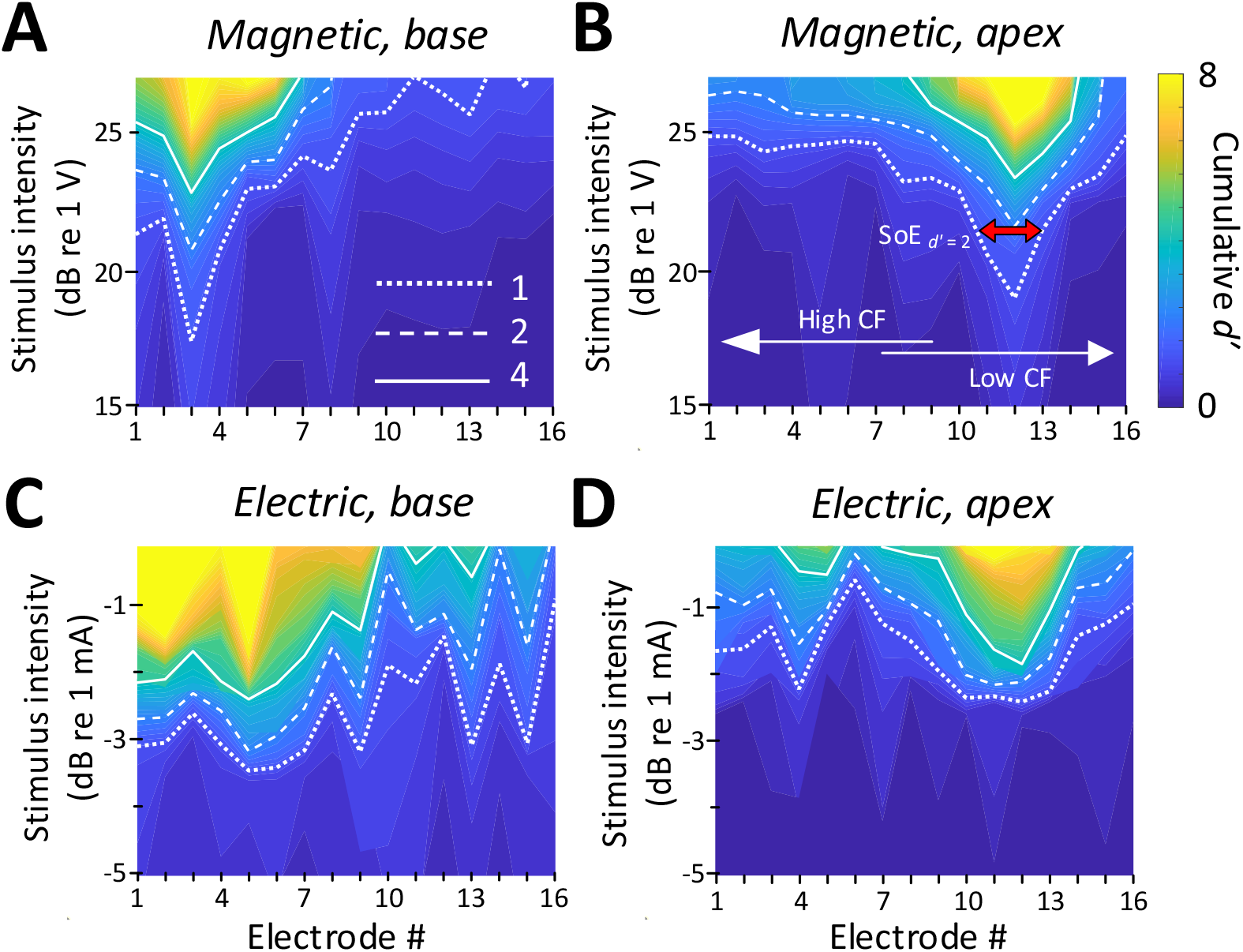
STCs in response to magnetic and electric stimulation. **(A and B)** Spatial tuning curves (STCs) in response to magnetic and electric stimulation delivered to the basal turn of the cochlea (aMUA signals quantified with d-prime analysis – see text). Dotted, dashed and solid lines are contours for cumulative *d’*-values of 1, 2, and 4. **(C and D)** STCs in response to magnetic and electric stimulation of the apical turn. The color bar on the right side of panel B applies to all panels. The red arrow in panel B indicates the spread of excitation (SoE – see text).

In contrast to the relatively narrow spectral spread of IC activity arising from magnetic stimulation, the spread from electric stimulation was considerably wider (Figs. 3C and D), consistent with findings in previous studies(14, 33-36). Nevertheless, BSs again showed evidence of tonotopic organization, i.e., stimulation of the basal turn was centered in the ventral portion of the IC while stimulation of the apex resulted in activation centered in the dorsal portion.

### Spatially confined responses elicited by magnetic stimulation

To quantify the spread of excitation across modalities, we measured the width of the *d′* = 1 trace (i.e., the distance between the ventral-most and dorsal-most electrodes exhibiting a supra-threshold response) at the stimulus amplitude for which the BS reached *d′* levels of 2 and 4 (the red arrow in Fig. 3B illustrates a sample calculation for a *d′* level of 2) (see MATERIALS AND METHODS). The spatial spread, i.e., the distance between the dorsal-most and ventral-most responding electrodes, was then converted into a spectral spread, i.e., the width of the corresponding frequency bands, derived from Fig. 1G (Figs. 4A and B).

**Fig. 4.**
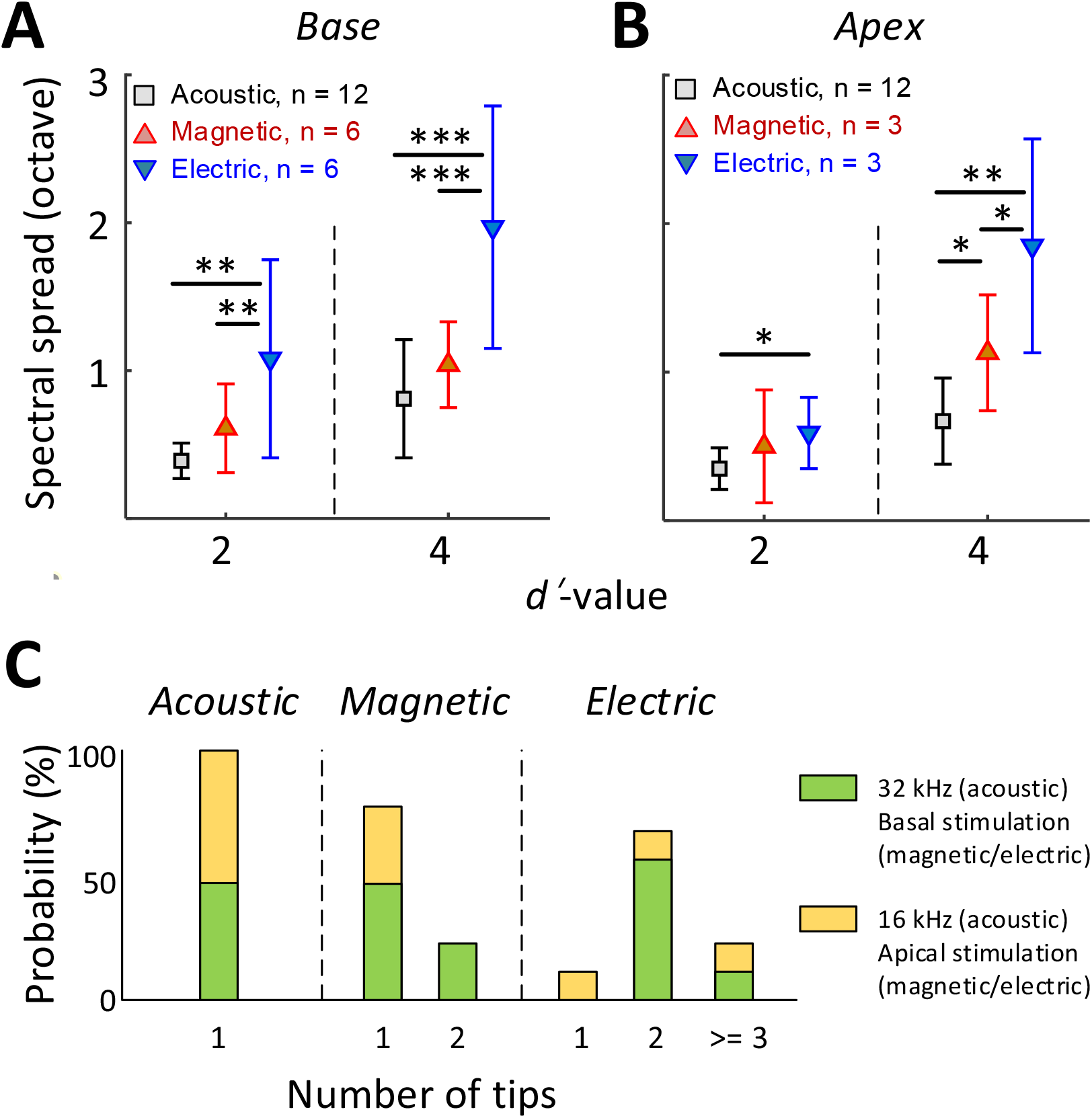
The spread of activation is narrower for magnetic vs. electric stimulation. **(A and B)** Mean ± SD for the spread of excitation by acoustic, magnetic, and electric stimulation delivered to the basal (A) or apical (B) turn of the cochlea (* *p* < 0.05, ** *p* < 0.01, *** *p* < 0.001) (**C)** The number of tips in the STCs for each stimulus modality (see text).

In all cases, i.e., for both basal and apical stimulation locations and both *d′* levels (moderate and high discrimination levels), the spread of activation from electric stimulation was wider than that from acoustic stimulation (p < 0.05; Fig. 4A and B). The spread from electric stimulation was also significantly wider than that from magnetic stimulation for all stimulation conditions, except for the specific case of apical stimulation at *d′* = 2. When the spread of magnetic stimulation was compared to that from auditory stimuli, there were no statistical differences for stimulation of the basal turn (both moderate and high discrimination levels) (p > 0.05) and the apical turn for the moderate discrimination level (*d′* = 2). The spread from magnetic stimulation was significantly wider than that from auditory stimulation only for apical stimulation and only for the high discrimination level (p = 0.031).

The number of peak ‘tips’ observed in the STCs differed across the stimulus modalities (Fig. 4C). For example, 8 of the 9 STC profiles generated in response to electric stimulation exhibited two or more tips leading to an average of 2.33 per profile. In contrast, 7 of the 9 STC profiles for magnetic stimulation had only a single peak tip, with the remaining two profiles showing double tips (average of 1.22). All profiles for auditory stimulation had a single peak only. The number of tips for electric stimulation was statistically higher than that from magnetic stimulation (p = 0.007) or from acoustic stimulation (p < 0.001). The difference between the number of tips for magnetic vs. acoustic stimulation was not statistically significant (p > 0.05). Taken together, these results indicate that the spread of activation was significantly narrower for magnetic vs. electric stimulation, regardless of the location at which stimulation was delivered. While much additional testing is required, the narrower spectral spread from magnetic stimulation suggests the possibility that a coil-based CI may create narrower and more independent spectral channels, thus offering the potential for improved performance of an implant.

### Larger dynamic range with magnetic stimulation

The rate-level functions to magnetic and electric stimulation (measured at the BS) are shown in Fig. 5A and B, respectively. The average DR across the population was smaller for magnetic stimulation (10.05 ± 4.18 dB V, n = 6; basal stimulation, measured at the BS) than for acoustic stimulation (25.96 ± 9.17 dB SPL; 32 kHz tone; p < 0.001) but larger than that for electric stimulation (3.24 ± 0.99 dB mA; p = 0.0031) (Fig. 5D), suggesting better discrimination resolution for stimulus intensity. The differences in DR did not arise from different neuronal response levels as the maximum response evoked by magnetic stimulation was comparable to that evoked by acoustic or electric stimulation, i.e., no statistically significant differences between any pair of modalities (Fig. 5C). Note that in some experiments with magnetic stimulation, response rates did not saturate, even at the maximum stimulation levels tested here (Fig. 5A), suggesting that the DRs reported here may be underestimated.

**Fig. 5.**
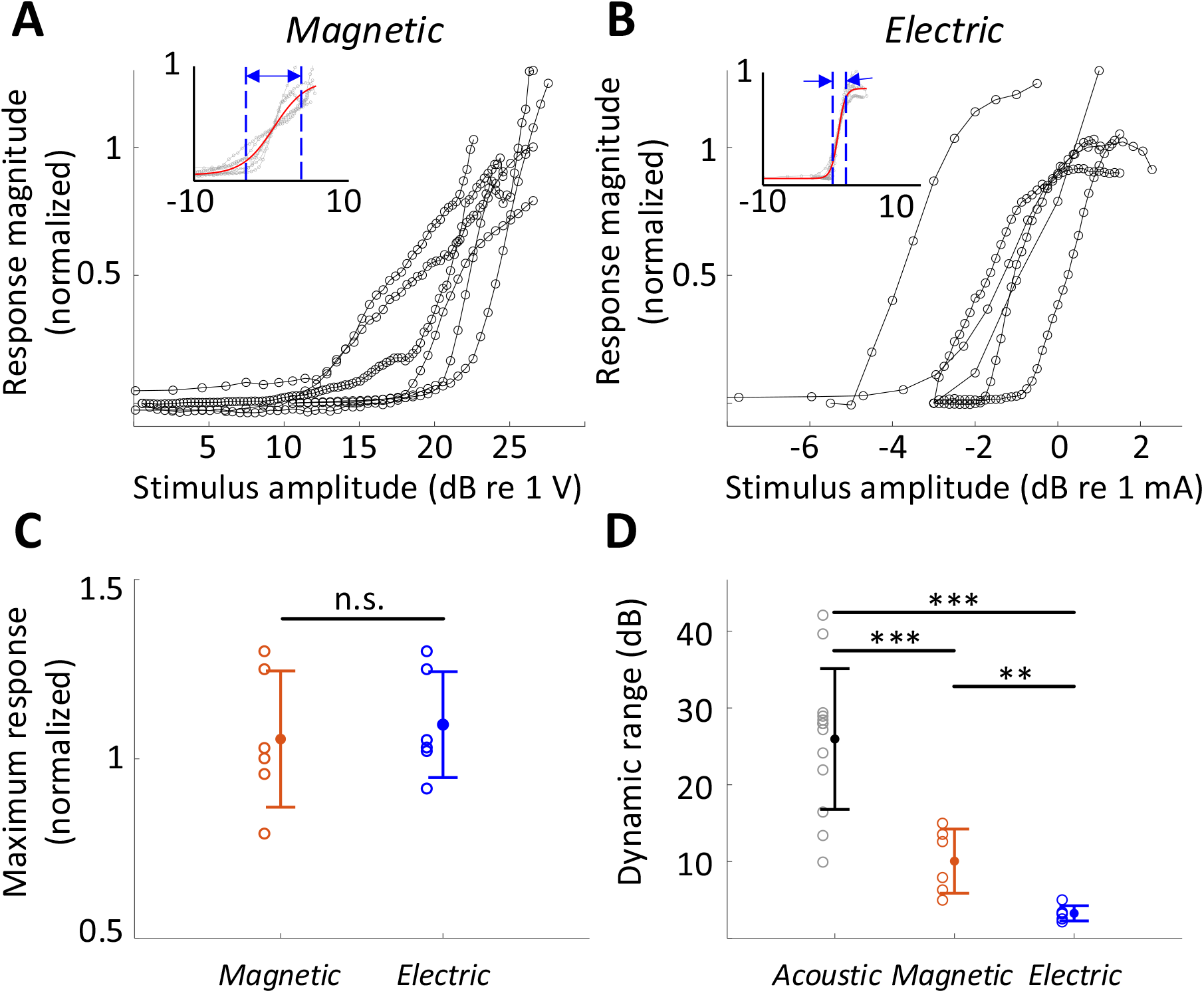
The dynamic range for magnetic stimulation is wider than that for electric stimulation. **(A and B)** Normalized responses rates as a function of stimulus intensity for magnetic (A), and electric (B) stimulation (basal turn). Each line is the averaged response curve from one animal. Insets show the same data normalized such that 50% of the peak response was assigned the level of 0 dB; the red line shows the best-fit curve to all raw data points. **(C)** Individual points are the distribution of the maximum response rates to magnetic and electric stimulation. Vertical lines show mean ± SD. Each response rate was normalized by the maximum response to acoustic stimulation obtained from the same animal. **(D)** Individual points are the distribution of dynamic ranges for each mode of stimulation. Vertical lines show mean ± SD (* *p* < 0.05, ** *p* < 0.01, *** *p* < 0.001).

### Responses to micromagnetic stimulation in hair cell ablated animals

As a final control experiment, we confirmed that magnetic and electric responses were similar in animals with chronically lesioned hair cells. These experiments used gentamicin to cause complete loss of inner and outer hair cells in the basal half of the cochlea (Fig. 6A)(37). After 10 days, hearing was evaluated with acoustic ABR and distortion product otoacoustic emissions (DPOAEs). DPOAEs were absent and ABR thresholds were beyond the range of the acoustic system or markedly increased (> 75 dB SPL, data not shown); small amounts of residual ABR can sometimes arise from acoustic cross-talk to the contralateral unlesioned ear(38). From these gentamicin-treated animals (n = 2), responses to electric and magnetic stimulation in the basal turn remained robust (Fig. 6B). Immunostaining in the basal turn at 32 kHz for hair cells (Myo7a, white) and actin-containing supporting structures (Phalloidin, red) showed no remaining inner and outer hair cells (Fig. 6A), eliminating the possibility that observed responses to magnetic or electric stimulation arose from inadvertent activation of hair cells.

**Fig. 6.**
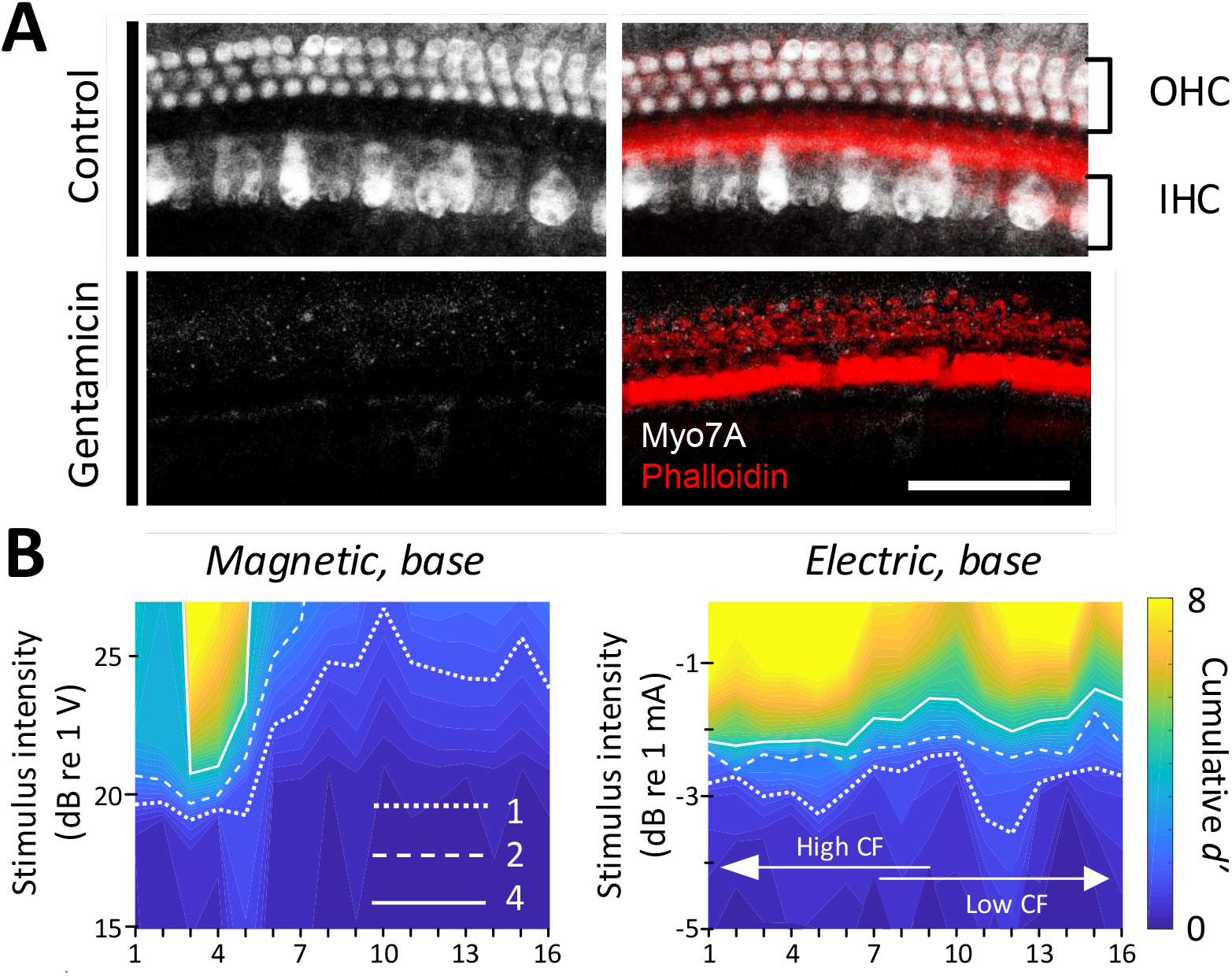
Responses in chronically deafened ears. **(A)** Gentamicin caused complete loss of inner and outer hair cells (white, Myo7A). Supporting structures of the organ of Corti were stained with Phalloidin (red). Scale bar: 50 µm. **(B)** STCs of the IC responses to magnetic and electric stimulation at the base measured in a gentamicin-treated mouse. Dotted, dashed, and solid lines are contours for cumulative *d’*-values of 1, 2, and 4, respectively.

## DISCUSSION

We used a combination of ABR measurements and multiunit recordings to demonstrate that magnetic stimulation, delivered from a bent-wire micro-coil inserted into the cochlea, can effectively drive the auditory pathways. Magnetic stimulation evoked multi-peaked ABRs, suggesting that coil-based stimulation was indeed capable of activating SGN processes that, in turn, led to activation of central nuclei in the auditory pathway. Multi-electrode recordings obtained in the IC showed that responses were robust, narrowly confined, and tonotopically organized. The responses from basal and apical micro-coil locations were narrow and showed little or no overlap between them, whereas responses from electric stimulation showed considerable overlap for stimulation at the same locations. The number of peak tips observed in the ST profiles generated from the IC recordings was also significantly lower for magnetic stimulation than for electric stimulation. suggesting the spread to encompass additional cochlear turns was less with magnetic stimulation. Taken together, the results strongly suggest that coil-based activation of the cochlea is more narrowly confined than that of electric stimulation.

A number of control experiments and experimental safeguards were used to verify that observed responses arose from magnetic stimulation of SGNs and not from other factors. First, the impedance from the coil to ground was monitored before and after each experiment and remained consistently above 200 MΩ, eliminating the possibility that observed responses arose from direct electrical activation, e.g., from leakage of the stimulus current into the surrounding perilymph. Second, the DC resistance across the coil leads was also monitored regularly and it too remained stable (∼10 Ω), eliminating the possibility that a broken coil might be activating neurons capacitively. Third, measurements of the temperature change produced by coils were previously shown to be less than 1° C(17, 20), greatly reducing the possibility that observed responses arose from some type of temperature shock. The fourth set of control experiments arose from concerns that observed responses could be mediated through activation of hair cells, e.g., micro-movements of the coil during the delivery of stimulus current could result in transmission of a pressure wave through the scala tympani(39). To eliminate this possibility, we injected DI water into the cochlea after completing the measurements of auditory responses. The resulting osmotic shock led to loss of responses to subsequent auditory stimuli, even at SPL levels of 75 dB, strongly suggesting a loss of hair cell functionality. To provide even stronger assurance, we injected gentamicin into the cochlea in a subset of animals; robust responses to magnetic and electrical stimulation persisted in these animals but not to acoustic stimulation. Post-mortem immunochemical staining of the cochlea revealed a complete absence of hair cells, eliminating the possibility that responses arising from micro-coils were mediated through some type of magneto-acoustic effect.

### Activation from magnetic stimulation is spatially confined

Previous studies have reported that the spread of current from electrodes inserted into the scala tympani recruits SGNs across spatially extensive regions, thereby limiting the spectral specificity of artificial sound encoding(14, 33-36). For example, a study in cat(14) compared the spectral spread arising from acoustic stimulation to that from monopolar and bipolar electric stimulation by similarly recording neural activity across the tonotopic axis of the IC. Although the spectral spread from bipolar electric stimulation was significantly narrower than that from monopolar stimulation in that study, both were substantially wider than that from acoustic stimulation (8.23- and 4.25-fold, respectively). Another study in mice(40) reported that spatial tuning curves (STCs) for electric stimulation were 3-4 times broader than those for pure tones. Thus, the wide spread of activation from electric stimulation measured here (3.14 times larger than that from monotones) is in good agreement with previous reports. Computational models that explore the reasons for the lack of confinement with electric stimulation find that the high conductivity of the perilymph in the scala tympani causes significant spread of the electric field arising from each electrode(41-44). In addition, because the SGN processes targeted by stimulation are situated on the other side of the high-resistance bony wall of the scala tympani, it is necessary to employ stronger stimulus amplitudes, thereby exacerbating the spread.

In contrast to the electric fields arising from electrodes, the spread of activation from coils measured here was significantly narrower. While our experiments do not identify the reasons for the lower spread, magnetic stimulation is known to have some intrinsic advantages over electrodes and electric stimulation. For example, the physics underlying the generation and spread of magnetic fields (Maxwell’s equations) ensures that the induced electric fields are confined to tight regions around the magnetic flux. In addition, the magnetic fields induced by the flow of electric current through coils are highly permeable to all biological materials and thus are relatively impervious to bone and other high-resistance materials within the cochlear environment. As a result, magnetic fields pass readily through the high-resistance bony wall of the scala tympani, without the need to increase stimulation amplitudes (thus, limiting additional spread). Although magnetic fields are not thought to activate neurons directly, time-varying magnetic fields induce electric fields, and therefore magnetic fields ‘carry’ the electric field across the walls of the scala tympani where they can induce activation of SGN processes.

### Improved dynamic range with magnetic stimulation over electric stimulation

In addition to the low spectral resolution associated with electric stimulation, the dynamic range for encoding sound intensity is also limited. For example, the dynamic range for listeners with normal hearing is approximately 120 dB while the dynamic range for CI users is typically restricted to 10–20 dB(45). The small dynamic range for electric stimulation is related in part to the wide spread of activation(46). SGN populations with similar characteristic frequencies encode sound intensity together which allows a wider dynamic range to be perceived at downstream auditory circuits(46-48); simultaneous activation of all such fibers with electric stimulation compresses the DR. The relatively small dynamic range for electric stimulation results in the need for amplitude compression of the acoustic signal with CIs(45). The practical implications of this compression are potential decreases in speech recognition(49, 50), particularly in the presence of increased background noise(51), as well as reductions in sound quality(52, 53).

Our measurements suggest that the dynamic range for magnetic stimulation from micro-coils was approximately 3 times greater than that for electrical stimulation. It is likely that the gradual recruitment of SGNs with magnetic stimulation contributed to the wider dynamic range. Further, the responses to magnetic stimulation were not saturated at the peak stimulus amplitudes we used here, suggesting the DR values reported here may be underestimated. The expanded DRs for magnetic stimulation provide additional support for the potential of micro-coil-based CIs to enhance the quality of CI-induced hearing.

### Future efforts and limitations of micro-coil stimulation of the cochlea

The ability to create narrow spectral channels with micro-coils, even in the tiny cochlea of the mouse, is encouraging as it suggests that coil-based stimulation has the potential to create a larger number of independent spectral channels in clinical use. Further, the relatively simple and compact bent-wire coils used here raise the possibility that CIs can be manufactured that are structurally similar to existing devices. This too is encouraging because such an approach raises fewer short- and long-term safety concerns. While much additional development and testing is needed, we believe that these findings clearly suggest that further investigation of coil-based stimulation of the cochlea is warranted.

Despite the encouraging results in this first assessment of coil-based stimulation of the cochlea, several key elements of micro-coil design and performance will need to be optimized before they can be considered for human trials. For example, the amplitude of the electric current that flowed through the coil was typically quite large, ∼770 mA at threshold, raising concerns about power consumption and battery life. While the relatively low impedance of micro-coils (typically ∼10 Ω) helps to reduce the I^2^ × R power consumption of coil-based devices and thus compensates somewhat for the high current levels, supplied electric energy for a single magnetic ‘pulse’ (52 µJ) is still considerably higher than that for a single electrical biphasic pulse (245 nJ – based on 700 µA and 10 kΩ impedance). It is likely that power consumption in future micro-coil devices can be significantly improved using a number of changes that are relatively straightforward to implement. For example, the coil design used here was identical to that used for stimulation of cortex; tailoring the coil design to optimize SGN activation could potentially reduce power consumption by an order of magnitude or more(19, 54). In addition, switching from the platinum-iridium wires used here to higher conductivity materials such as silver or gold could reduce power consumption in half. Finally, the incorporation of magnetic cores into the coil could potentially reduce thresholds by several orders of magnitude. Importantly, future testing will need to take place in animal models whose cochleae better resemble those of humans. Larger size scala tympani associated with such animals will likely require stronger activation thresholds to compensate for the increased distance to targeted neurons, although they will also allow larger wire sizes, possibly offsetting the difference. The high stimulus amplitudes associated with coils also raise concerns about electrical safety, although it is important to remember that the flow of electrical current through the coil is electrically isolated from the surrounding tissue. Advanced control circuits can be incorporated into future designs to further minimize the potential for tissue damage. Even without the electrical concerns, the high current levels required to activate SGNs may produce temperature changes that could exceed safe limits; these too will need to be evaluated prior to chronic use of coil-based implants.

Finally, it will also be necessary to build and test devices that incorporate multiple coils to ensure that they can indeed be reliably developed and safely implanted as well as to determine the minimum separation for which coils remain independent. The ability to match the structure and mechanical properties of coil-based CIs will be highly beneficial as it will allow existing fabrication techniques and surgical procedures to be harnessed, thereby facilitating the transition into clinical practice.

## MATERIALS AND METHODS

### Animal preparation

All procedures were approved by the Institutional Animal Care and Use Committee of Massachusetts Eye and Ear, and carried out in accordance with the NIH Guide for the Care and Use of Laboratory Animals. CBA/CaJ mice were purchased from Jackson Laboratories or bred in house. Mice of either sex aged 6 – 16 weeks were used in the experiments. Experiments were conducted in an acoustically and electrically isolated walk-in chamber kept at 32 – 36 °C. Mice were anesthetized for the duration of experiments with ketamine (100 mg/kg) and xylazine (10 mg/kg). Animals’ anesthesia level and heart rate were regularly monitored, and one third of the initial ketamine/xylazine dose (i.e. 33 mg/kg and 3 mg/kg, respectively) was given as needed.

The left cochlea was accessed surgically. A postauricular incision was made and the underlying tissue and musculature were dissected to expose the bulla. A bullotomy was performed by carefully rotating a 28 G needle and enlarging the hole with fine forceps to expose the round window. The left pinna with skin and tissue extending into the external auditory canal was cut and removed to expose the tympanic membrane. To access the contralateral inferior colliculus (IC), the postauricular incision was extended to over midline, and a craniotomy was made just caudal to the temporo-parietal suture and to the contralateral side of the midline with a scalpel.

### Stimulation

All stimuli were generated using LabVIEW and MATLAB software controlling custom-made system based on National Instruments 24-bits digital input/output boards.

#### Acoustic stimulation

A custom acoustic system coupled to a probe tube was inserted into the external ear canal close to the tympanic membrane, with two miniature earphones (CDMG150 008-03A, CUI) serving as sound sources. Acoustic stimuli were pure tone pips of 5 ms duration.

#### Magnetic and electric stimulation

Magnetic stimulation was presented using custom-made micro-coils (MicroProbes, Gaithersburg, MD), which are highly similar to ones used in previous studies for stimulation of other, non-cochlear regions of the CNS(17, 20). The coil was fabricated by bending a 25 µm-diameter platinum-iridium into a U-shape (Fig. 1C); The length of the coil was 3 mm and the width was 175 µm. The direct current (DC) resistance of the coil was in the range of 8-10 Ω. The coil wire was coated with 5-µm-thick parylene for electrical insulation. Coils were tested before and after each experiment to ensure that there was no inadvertent leak of electric current from the coil to the cochlea. To this end, each coil was submerged in NaCl solution (0.9%) and the electric resistance between one of the coil’s terminal ends and an electrode immersed in the solution was measured; resistances above 200 MΩ were considered sufficient for insulation. At least three individual coils with an identical design were tested. To deliver magnetic stimulation, a micro-coil was inserted through the round window, and responses to a range of stimulation parameters delivered to the basal turn were captured. After completion of experiments at the basal turn of the cochlea, a cochleostomy was performed near the apex, allowing analogous experiments to be performed at the apical turn as well. The stimulus was generated by a function generator based on National Instruments 24-bits digital input/output boards and amplified by a voltage amplifier with a gain of 9 V/V and a bandwidth of 70 kHz (PB717X, Pyramid Inc. Brooklyn, NY, USA). The voltage amplifier was powered by a commercial battery (LC-R1233P, Panasonic Corp., Newark, NJ, USA). The stimulus waveform was a positive-going ramp with a rise time of 25 µs. The fall time was set to 0 μs but was limited by the sampling rate of the hardware (100 kHz). The amplitude of the waveform from the function generator was 0 V to 1.7 V. The output of the amplifier was 0 V to 15.3 V. The peak levels of magnetic stimulation are limited by Joule heating of the small wires that comprise the micro-coils and the resulting potential to induced failure. At a given stimulus strength, the waveform was presented a minimum of 39 times with a pulse rate of 25 pulses/s.

Once completed, the coil was removed and replaced with a micro-electrode so that an analogous series of electric stimulation experiments could be performed at the same location. Electric stimulation was delivered in a monopolar configuration. The stimulating electrode was a conical platinum-iridium tip with a resistance of 10 kΩ, a height of ∼125 µm and a base diameter of 30 µm (Microprobes for Life Science, Gaithersburg, MD, USA; PI2PT30.01 A10). An EMG needle was inserted into the neck muscle to serve as a return electrode. At least three individual electrodes with an identical design were tested. Like micro-coils, the stimulating electrodes were inserted through the round window (for basal stimulation) or via cochleostomy in the apical turn for intracochlear stimulation. Electric stimuli were also controlled by the same function generator used for magnetic stimulation. Stimulus waveforms were rectangular biphasic pulses with phase duration of 25 µs and no inter-phase interval. Stimulation amplitudes ranged from 0 µA to 1,000 µA. The stimulus for each amplitude was repeated at least 39 times with a pulse rate of 25 pulses/s.

Magnetic stimulation responses were assayed first in some animals, whereas electric stimulation responses were assayed first in other animals. The same placements were used for either the micro-coils or the electrodes. We terminated the experiments once the elicited ABR or IC responses to magnetic stimulation were no longer robust. A complete set of basal and apical stimulations were tested in 2 animals, a complete set of only basal stimulation was tested in 4 additional animals, and a complete set of only apical stimulation was tested in 1 additional animal.

### Data acquisition and analysis

#### DPOAE and ABR

DPOAE and ABR were recorded as previously described(55). A custom acoustic system was inserted into the external ear canal close to the tympanic membrane. DPOAEs were measured as ear canal pressure in response to two tones presented into the ear canal (*f*_*1*_ and *f*_*2*_, with *f*_*2*_ / *f*_*1*_ = 1.2 and *f*_*1*_ being 10 dB above *f*_*2*_) at half-octave steps from *f*_*2*_ = 5.66 – 45.25 kHz, and in 5 dB intensity increments from 10 to 80 dB SPL. DPOAE thresholds were defined as the *f*_*2*_ intensity required to generate a DP response of 10 dB SPL over noise floor. ABR responses to 5 ms tone pips were measured between subdermal electrodes (adjacent to the ipsilateral incision, at the vertex, and near the tail), amplified 10,000 times and filtered through a 0.3–3.0 kHz band-pass filter. For each frequency, the sound level starting below the threshold was increased in 5 dB-steps and 512 responses. ABR thresholds were defined as the lowest level at which a repeatable waveform could be visually detected.

#### Inferior Colliculus recordings

Neural activity in the IC was recorded using a 16-channel, single-shank electrode array (177 μm^2^/site, center-to-center electrode spacing of 50 μm; NeuroNexus Technologies, Ann Arbor, MI; A1×16-3mm-50-177). Recordings were collected at a sampling rate of 25 kHz and analyzed offline using custom-written MATLAB scripts. While multi-unit activities (MUAs) were evident in all channels (Fig. S1), distinguishable action potentials could be detected from only a few channels, typically less than 3. Thus, data analysis was based on MUAs, which were quantified by an analog representation of multi-unit activity (aMUA). aMUA reflects the voltage signal power within the frequency range occupied by action potentials (56-60). This approach is advantageous over the more traditional measure of MUA based on thresholding and spike-detection since aMUAs are not biased by free parameters (e.g., threshold levels), and provide a high signal to noise ratio(59). aMUA was measured as follows: (1) To extract MUA, the raw recordings were band-pass filtered between 325 and 6000 Hz (Butterworth IIR); During this process, low-frequency local field potentials and high-frequency electric noise signals were removed (Fig. S1A). (2) the recorded signals prior to 2 ms after stimulus onset were excluded from subsequent analyses due to stimulus artifact (Fig. S1B). (3) The extracted MUA signal was then rectified to take the absolute value of response magnitude over time. In addition, to reduce aliasing artifact, the processed signal was next low pass filtered at 475 Hz (Butterworth IIR), and downsampled from 25 kHz to 12 kHz (Fig. S1C). (4) The area under the curve for the time period 2 – 15 ms following stimulus onset was then calculated.

#### d-prime analysis

To quantify the change in neural responses to stimulus intensity, the discrimination index, *d′* (d-prime) was calculated by comparing aMUAs across successive pairs of stimulus levels in each electrode(14, 61). Based on aMUAs to a given stimulus level and those to the next higher level, a *d′* value with unequal variance was calculated as

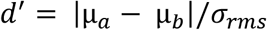

where µ and σ_*rms*_ are the mean aMUA and common standard deviation, respectively(62, 63). The *d′* values were then accumulated up to each stimulus intensity to calculate the cumulative *d′*.

#### Construction of spatial tuning curve (STC)

STCs were used to compare spectral spreads from acoustic, magnetic and electric stimulation. Based on the cumulative *d′* values, an n × m matrix was constructed, where n corresponded to stimulus intensity and m to the electrode number. Iso-*d′*-contour-lines were derived by interpolating the matrix using MATLAB software. In all STCs (Fig. 1D-F, Fig. 3A-D), contours for cumulative *d′* levels of 1, 2, and 4 are shown. The stimulation threshold was selected as the cumulative *d′* value of 1, and the best site (BS) was determined from the minima of the *d′* = 1 iso-contour. From the acoustic stimulation, the relationship between tone frequency and BS was plotted and used to interpolate the characteristic frequency of each electrode.

The spatial spread of the response was calculated as the distance between the ventral-most and dorsal-most electrodes exhibiting a supra-threshold response (*d′* > 1). Using the characteristic frequency of each electrode obtained from the acoustic stimulation, the spatial spread could then be converted to the spectral spread of cochlear excitation, i.e., the width of the corresponding frequency bands.

The spread of excitation was also evaluated by measuring the number of peaks observed in the ST profiles. The number of peaks was measured by the number of isolated electrode groups exhibiting a supra-threshold response (*d′* > 1) at the stimulus intensity that elicited a cumulative *d′* value of 2.

#### Statistical evaluation

All data were presented as the mean ± standard deviation. A student t-test was performed to determine whether the difference between data was significant.

### Deafening

After recording acoustic responses, animals were acutely deafened by gently infusing 5 µl of distilled water through the round window to cause osmotic stress(28). Successful deafening was confirmed 10 minutes after injection by re-measuring auditory brainstem response (ABR) to acoustic stimuli – a sharp increase in ABR thresholds (≥ 75 dB SPL) or complete elimination of the waveform was considered evidence that hair cells were no longer functioning. Small amounts of residual ABR were attributed to acoustic cross-over to the contralateral, non-deafened ear (Harrison et al, 2013).

To chronically deafen animals through lesions of hair cells, we used the ototoxic aminoglycoside antibiotic gentamicin. The round window niche was exposed in anesthetized animals, and a small piece of gelfoam sponge soaked in 200 µg gentamicin in distilled water was applied to the round window membrane(37). The skin was closed with sutures. The animal received post-surgical analgesia with meloxicam (2 mg/kg) and buprenorphine (0.05 mg/kg).

Ten days after this procedure, the absence (or near absence) of ABR and DPOAE responses were used to confirm successful deafening; these animals were then used for electric and magnetic stimulation experiments.

### Cochlear whole mounts and confocal fluorescence immunohistochemistry

After completion of stimulation procedures, deeply anesthetized animals were intracardially perfused with 4% paraformaldehyde (PFA), both cochleae were extracted and processed as previously described(55). PFA was gently perfused through the round and oval windows. Cochleae were post-fixed for 2 hours in 4% PFA and decalcified in 0.12 M (ethylenediaminetetraacetic acid) EDTA for 72 hours. The decalcified spiraling cochleae were microdissected into 4 – 6 pieces, blocked with 5% normal horse serum (NHS) and 0.3% Triton X-100 (TX-100) in PBS for 30 min at room temperature, and immunostained to label hair cells overnight at room temperature with rabbit anti-myosin 7A (1:200, #25-6790 Proteus Biosciences, Ramona, CA) diluted in 1% normal horse serum with 0.3% TX. After washing in PBS, cochlear pieces were incubated with Alexa Fluor 488-conjugated goat anti-rabbit antibody (#A-11008) and Alexa Fluor 647-conjugated phalloidin (#A22287) at 1:200 (Invitrogen, Carlsbad, CA) for 90 min. A cochlear frequency map was created by applying a custom ImageJ plug-in (https://www.masseyeandear.org/research/otolaryngology/eaton-peabody-laboratories/histology-core) to images acquired at low magnification (10x objective) on a fluorescent microscope (E800, Nikon, Melville, NY). Cochlear whole-mounts were subsequently imaged with a confocal microscope (SP8, Leica, Wetzlar, Germany) using a 63x glycerol-immersion objective (1.3 N.A) at the 32 kHz cochlear frequency region.

## Supplementary Materials

Fig. S1. IC responses to acoustic, magnetic and electric stimulation

## Acknowledgments

We thank Kenneth Hancock for his expert technical assistance.

---

**Fig. S1.**
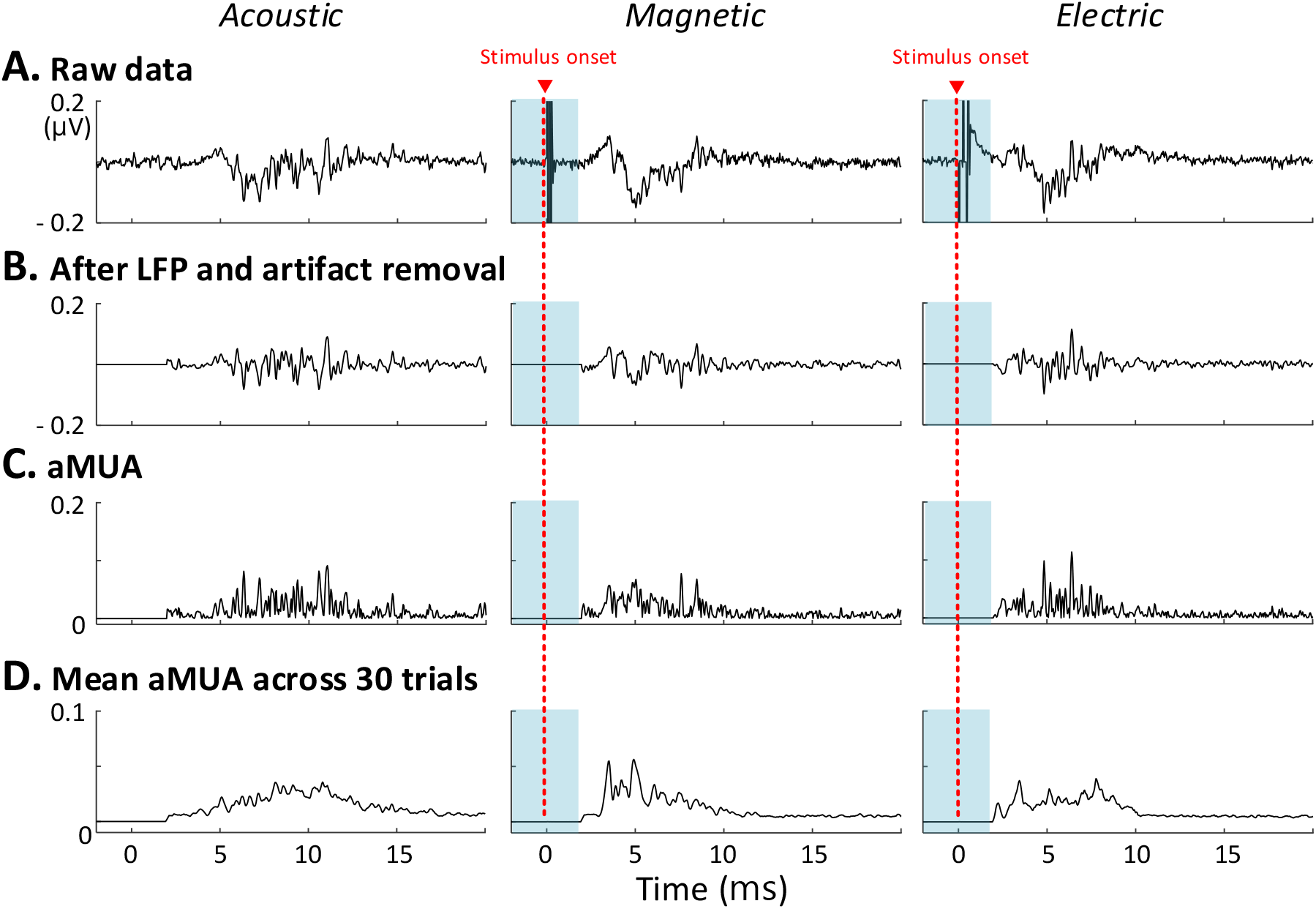
IC responses to acoustic, magnetic and electric stimulation. **(A)** Representative raw recordings of single traces of the neural activity evoked by each stimulation modality were captured by the same recording electrode (positioned in IC). **(B)** Local field potential were removed from the raw trace by band-pass filtering. Stmulus artifact was removed prior to 2 ms after stimulus onset from the magnetic and electric stimulation responses by setting the response to zero. **(C)** Analog representation of multi-unit activity (aMUA; see Methods) **(D)** Mean aMUA across 30 trials

## References

1. WHO. Deafness and hearing loss. 2021.

2. Gurgel RK, Ward PD, Schwartz S, Norton MC, Foster NL, Tschanz JT. Relationship of hearing loss and dementia: a prospective, population-based study. Otol Neurotol. 2014;35(5):775.

3. Firszt JB, Holden LK, Skinner MW, Tobey EA, Peterson A, Gaggl W, et al. Recognition of speech presented at soft to loud levels by adult cochlear implant recipients of three cochlear implant systems. Ear Hear. 2004;25(4):375–87.

4. Holden LK, Finley CC, Firszt JB, Holden TA, Brenner C, Potts LG, et al. Factors affecting open-set word recognition in adults with cochlear implants. Ear Hear. 2013;34(3):342–60.

5. Moberly AC, Lowenstein JH, Nittrouer S. Word Recognition Variability With Cochlear Implants: “Perceptual Attention” Versus “Auditory Sensitivity”. Ear Hear. 2016;37(1):14–26.

6. Moberly AC, Lowenstein JH, Nittrouer S. Word Recognition Variability With Cochlear Implants: The Degradation of Phonemic Sensitivity. Otol Neurotol. 2016;37(5):470–7.

7. Friesen LM, Shannon RV, Baskent D, Wang X. Speech recognition in noise as a function of the number of spectral channels: Comparison of acoustic hearing and cochlear implants. J Acoust Soc Am. 2001;110(2):1150–63.

8. Jiam NT, Caldwell MT, Limb CJ. What does music sound like for a cochlear implant user? Otol Neurotol. 2017;38(8):e240–e7.

9. Fishman KE, Shannon RV, Slattery WH. Speech recognition as a function of the number of electrodes used in the SPEAK cochlear implant speech processor. J Speech Lang Hear Res. 1997;40(5):1201–15.

10. Fu Q-J, Nogaki G. Noise susceptibility of cochlear implant users: the role of spectral resolution and smearing. J Assoc Res. 2005;6(1):19–27.

11. Carlyon RP, Long CJ, Deeks JM, McKay CM. Concurrent sound segregation in electric and acoustic hearing. J Assoc Res. 2007;8(1):119–33.

12. Hernandez VH, Gehrt A, Reuter K, Jing Z, Jeschke M, Schulz AM, et al. Optogenetic stimulation of the auditory pathway. J Clin Investig. 2014;124(3):1114–29.

13. Dieter A, Duque-Afonso CJ, Rankovic V, Jeschke M, Moser T. Near physiological spectral selectivity of cochlear optogenetics. Nat Commun. 2019;10(1):1–10.

14. Middlebrooks JC, Snyder RL. Auditory prosthesis with a penetrating nerve array. J Assoc Res Otolaryngol. 2007;8(2):258–79.

15. Middlebrooks JC, Snyder RL. Intraneural stimulation for auditory prosthesis: modiolar trunk and intracranial stimulation sites. Hear Res. 2008;242(1-2):52–63.

16. Middlebrooks JC, Snyder RL. Selective electrical stimulation of the auditory nerve activates a pathway specialized for high temporal acuity. J Neurosci. 2010;30(5):1937–46.

17. Lee SW, Fallegger F, Casse BDF, Fried SI. Implantable microcoils for intracortical magnetic stimulation. Sci Adv. 2016;2.

18. Lee SW, Fried SI. Enhanced Control of Cortical Pyramidal Neurons with Micromagnetic Stimulation. IEEE Trans Neural Syst Rehabil Eng. 2017;25:1375–86.

19. Lee SW, Thyagarajan K, Fried SI. Micro-Coil Design Influences the Spatial Extent of Responses to Intracortical Magnetic Stimulation. IEEE Trans Biomed Eng. 2019;66:1680–94.

20. Ryu SB, Paulk AC, Yang JC, Ganji M, Dayeh SA, Cash SS, et al. Spatially confined responses of mouse visual cortex to intracortical magnetic stimulation from micro-coils. J Neural Eng. 2020;17.

21. Osanai H, Minusa S, Tateno T. Micro-coil-induced inhomogeneous electric field produces sound-driven-like neural responses in microcircuits of the mouse auditory cortex in vivo. Neurosci. 2018;371:346–70.

22. Minusa S, Osanai H, Tateno T. Micromagnetic stimulation of the mouse auditory cortex in vivo using an implantable solenoid system. IEEE Trans Biomed Eng. 2018;65:1301–10.

23. Mukesh S, Blake DT, McKinnon BJ, Bhatti PT. Modeling Intracochlear Magnetic Stimulation: A Finite-Element Analysis. IEEE Trans Neural Syst Rehabil Eng. 2017;25:1353–62.

24. Stiebler I, Ehret G. Inferior colliculus of the house mouse. I. A quantitative study of tonotopic organization, frequency representation, and tone-threshold distribution. J Comp Neurol. 1985;238:65–76.

25. Köppl C, Yates G. Coding of sound pressure level in the barn owl’s auditory nerve. J Neurosci. 1999;19:9674–86.

26. Nizami L. Estimating auditory neuronal dynamic range using a fitted function. Hear Res. 2002;167:13–27.

27. Shivdasani MN, Mauger SJ, Rathbone GD, Paolini AG. Inferior colliculus responses to multichannel microstimulation of the ventral cochlear nucleus: Implications for auditory brain stem implants. J Neurophysiol. 2008;99:1–13.

28. Chung Y, Hancock KE, Delgutte B. Neural coding of interaural time differences with bilateral cochlear implants in unanesthetized rabbits. J Neurosci. 2016;36:5520–31.

29. Melcher JR, Guinan JJ, Knudson IM, Kiang NYS. Generators of the brainstem auditory evoked potential in cat. II. Correlating lesion sites with waveform changes. Hear Res. 1996;93:28–51.

30. Biacabe B, Chevallier JM, Avan P, Bonfils P. Functional anatomy of auditory brainstem nuclei: Application to the anatomical basis of brainstem auditory evoked potentials. Auris Nasus Larynx. 2001;28:85–94.

31. Navntoft CA, Marozeau J, Barkat TR. Cochlear implant surgery and electrically-evoked auditory brainstem response recordings in C57BL/6 mice. J Vis Exp. 2019(143):e58073.

32. Miller CA, Woodruff KE, Pfingst BE. Functional responses from guinea pigs with cochlear implants. I. Electrophysiological and psychophysical measures. Hear Res. 1995;92(1-2):85–99.

33. Shannon RV. Multichannel electrical stimulation of the auditory nerve in man. II. Channel interaction. Hear Res. 1983;12(1):1–16.

34. O’Leary SJ, Black RC, Clark GM. Current distributions in the cat cochlea: a modelling and electrophysiological study. Hear Res. 1985;18(3):273–81.

35. O’Leary SJ, Richardson RR, McDermott HJ. Principles of design and biological approaches for improving the selectivity of cochlear implant electrodes. J Neural Eng. 2009;6(5):055002.

36. Micco AG, Richter CP. Tissue resistivities determine the current flow in the cochlea. Curr Opin Otolaryngol Head Neck Surg. 2006;14:352–5.

37. Heydt JL, Cunningham LL, Rubel EW, Coltrera MD. Round window gentamicin application: An inner ear hair cell damage protocol for the mouse. Hear Res. 2004;192:65–74.

38. Harrison AL, Negandhi J, Allemang C, D’Alessandro L, Harrison RV. Acoustic cross-over between the ears in mice (mus musculus) determined using a novel ABR based bio-assay. Can Acoust. 2013;41:5–10.

39. Kallweit N, Baumhoff P, Krueger A, Tinne N, Kral A, Ripken T, et al. Optoacoustic effect is responsible for laser-induced cochlear responses. Sci Rep. 2016;6:1–10.

40. Thompson AC, Wise AK, Hart WL, Needham K, Fallon JB, Gunewardene N, et al. Hybrid optogenetic and electrical stimulation for greater spatial resolution and temporal fidelity of cochlear activation. J Neural Eng. 2020;17.

41. Vanpoucke F, Zarowski A, Casselman J, Frijns J, Peeters S. The facial nerve canal: an important cochlear conduction path revealed by Clarion electrical field imaging. Otol Neurotol. 2004;25(3):282–9.

42. Briaire JJ, Frijns JH. Field patterns in a 3D tapered spiral model of the electrically stimulated cochlea. Hear Res. 2000;148(1-2):18–30.

43. Frijns JH, Kalkman RK, Briaire JJ. Stimulation of the facial nerve by intracochlear electrodes in otosclerosis: a computer modeling study. Otol Neurotol. 2009;30(8):1168–74.

44. Jolly CN, Spelman FA, Clopton BM. Quadrupolar stimulation for Cochlear prostheses: modeling and experimental data. IEEE Trans Biomed Eng. 1996;43(8):857–65.

45. Hong RS, Rubinstein JT, Wehner D, Horn D. Dynamic range enhancement for cochlear implants. Otology and Neurotology. 2003;24:590–5.

46. Viemeister NF. Intensity coding and the dynamic range problem. Hear Res. 1988;34:267–74.

47. Sachs MB, Abbas PJ. Rate versus level functions for auditory-nerve fibers in cats: tone-burst stimuli. The Journal of the Acoustical Society of America. 1974;56(6):1835–47.

48. Hudspeth AJ. Integrating the active process of hair cells with cochlear function. Nat Rev Neurosci. 2014;15(9):600–14.

49. Fu QJ, Shannon RV. Effects of dynamic range and amplitude mapping on phoneme recognition in nucleus-22 cochlear implant users. Ear Hear. 2000;21:227–35.

50. Loizou PC, Poroy O, Dorman M. The effect of parametric variations of cochlear implant processors on speech understanding. J Acoust Soc Am. 2000;108(2):790–802.

51. Zeng FG, Galvin JJ, 3rd. Amplitude mapping and phoneme recognition in cochlear implant listeners. Ear Hear. 1999;20(1):60–74.

52. Boike KT, Souza PE. Effect of compression ratio on speech recognition and speech-quality ratings with wide dynamic range compression amplification. J Speech Lang Hear Res. 2000;43(2):456–68.

53. Neuman AC, Bakke MH, Mackersie C, Hellman S, Levitt H. The effect of compression ratio and release time on the categorical rating of sound quality. J Acoust Soc Am. 1998;103(5 Pt 1):2273-81.

54. Frijns JHM, Briaire JJ, Grote JJ. The importance of human cochlear anatomy for the results of modiolus-hugging multichannel cochlear implants. Otol Neurotol. 2001;22:340–9.

55. Seist R, Tong M, Landegger LD, Vasilijic S, Hyakusoku H, Katsumi S, et al. Regeneration of Cochlear Synapses by Systemic Administration of a Bisphosphonate. Front Mol Neurosci. 2020;13:87.

56. Chung SH, Jones LC, Hammond BJ, King MC, Evans RJ, Knott C, et al. Signal processing technique to extract neuronal activity from noise. J Neurosci Methods. 1987;19(2):125–39.

57. King AJ, Carlile S. Responses of neurons in the ferret superior colliculus to the spatial location of tonal stimuli. Hear Res. 1994;81(1-2):137–49.

58. Choi Y-S, Koenig MA, Jia X, Thakor NV. Quantifying time-varying multiunit neural activity using entropy-based measures. IEEE Trans Biomed Eng. 2010;57(11):2771–7.

59. Schnupp JWH, Garcia-Lazaro JA, Lesica NA. Periodotopy in the gerbil inferior colliculus: Local clustering rather than a gradient map. Front Neural Circuits. 2015;9:1–21.

60. Sadeghi M, Zhai X, Stevenson IH, Escabí MA. A neural ensemble correlation code for sound category identification. PLoS Biol. 2019;17:1–41.

61. Macmillan NA, Creelman CD. Detection theory: A user’s guide: Psychology press; 2004.

62. Egan JP, Clarke FR. Psychophysics and signal detection. Technical Report, Bloomington, Indiana: Indiana University Hearing and Communication Laboratory; 1962.

63. Das A, Geisler WS. A method to integrate and classify normal distributions. Journal of Vision. 2021;21(10):1.

